# A programmable DNA-origami platform for studying protein-mediated lipid transfer between bilayers

**DOI:** 10.1101/610212

**Authors:** Xin Bian, Zhao Zhang, Pietro De Camilli, Chenxiang Lin

## Abstract

Non-vesicular lipid transport between bilayers at membrane contact sites plays important physiological roles. Mechanistic insight into the action of lipid transport proteins localized at these sites (bridge/tunnel versus shuttle models) requires a determination of the distance between bilayers at which this transport can occur. Here, we developed DNA-origami nanostructures to organize size-defined liposomes at precise distances and used them to study lipid transfer by the SMP domain of E-Syt1. Pairs of DNA ring-templated donor and acceptor liposomes were docked through DNA pillars, which determined their distance. The SMP domain was anchored to donor liposomes via an unstructured linker and lipid transfer was assessed via a FRET-based assay. We show that lipid transfer can occur over distances that exceed the length of SMP dimer, compatible with a shuttle model. The DNA nanostructures developed here can be adapted to study other processes occurring where two membranes are closely apposed to each other.

## Introduction

In eukaryotic cells, close appositions between the membranes of two different organelles mediated by protein tethers play a variety of functions in the control of organelle homeostasis, including lipid transport between the two adjacent bilayers via lipid-transport modules^1–4^. Examples of such modules are SMP (Synaptotagmin-like Mitochondrial lipid binding Protein) domains, which represent a branch within the TULIP (TUbular LIPid-binding proteins) domain superfamily^5–13^. SMP domains are found in proteins that function as tethers between the endoplasmic reticulum (ER) and other membranes^12, 14–17^, such as the three extended synaptotagmins (E-Syt1, E-Syt2 and E-Syt3), named tricalbin in yeast^18–20^. The E-Syts/tricalbins are anchored to the ER membrane via N-terminal hydrophobic hairpins, which are followed by the SMP domain and by variable numbers of C2 domains that mediate contacts of the ER membrane with the plasma membrane *in trans*^20–23^.

The SMP domain of the E-Syts dimerizes in an anti-parallel fashion to form a 90-Å elongated structure comprising a deep groove lined with hydrophobic residues that run from one end to the other of the module. This groove harbors glycerophospholipids, without selectivity for specific head groups^10^. A role of the SMP domain of the E-Syts in glycerolipid transport between bilayers has been supported by cell-free liposomes- and FRET-based assays and by evidence for a Ca^2+^ dependent role of E-Syt1 in the control of plasma membrane lipid homeostasis in response to acute perturbations^24–26^. However, how the SMP domain transfers lipids between two membranes remains unclear. Two models have been proposed^10^. In one model, the SMP domain directly bridges the two bilayers so that lipids can enter and exit it at its tips, and slide along the hydrophobic channel (tunnel model). In the second model, the SMP domain shuttles between the two bilayers, extracting lipids from one and delivering them to the other (shuttle model).

The tunnel model implies that transport can only occur when the two bilayers are separated by a distance that does not exceed the length of the SMP dimer (9 nm). The shuttle model, in contrast, is compatible with greatest distances, as long as such distance does not exceed the length of the predicted unstructured linker that connects the SMP domain to the bilayer of the ER. Analysis of ER-plasma membrane contact sites by cryo-electron tomography (cryo-ET) in mammalian cells showed that the ER-plasms membrane distance at contact sites can be variable and in the 10-30 nm range^22^, which seems too far for the dimeric SMP domain to span it. However, small, puncta-like closer appositions of the two bilayers (<10 nm), which would be compatible for the tunnel model, can occasionally be seen^22, 27^. Moreover, direct proof that E-Syts’ SMP domains can transfer lipids over a distance greater than their length is still lacking. Commonly used *in vitro* lipid transfer assays do not allow to control the distance of the two membrane bilayers. The goal of this study was to develop a method to assess the property of the SMP domain of the E-Syts, or of any other lipid-transport proteins or modules, to transfer lipids between bilayers over well-defined distances.

DNA origami is a technique that allows generating self-assembled and programmable DNA nanostructures^28–30^. Recently, it was reported that rationally designed DNA structures represent a versatile toolbox to grow, mold and fuse liposomes in a precise and deterministic way^31–38^. Here we have harnessed the engineering power of DNA in nanoscale to develop a distance-dependent lipid-transfer assay using DNA-origami-organized liposomes. The DNA nanostructure consists of two DNA rings that guide the formation of liposomes and a DNA pillar that keeps the two rings, and thus the two liposomes, at a predefined distance. This methodology offers a powerful solution to control spacing of membrane bilayers at a nanoscale distance and holds great potential for creating a generic platform for studies of protein-mediated lipid transfer or of other protein functions at sites where two membranes are closely apposed to each other. Our data provide evidence that SMP domains can transfer lipids over distances that exceed their length, compatible with the shuttle model of lipid transfer.

## Results

### A DNA-origami nanostructure to control the distance between lipid bilayers

In the commonly used liposome-based assays of protein-mediated lipid transport, the protein of interest is added to a suspension of donor and acceptor liposomes. In most cases, however, efficient transport requires tethering of the two bilayers, mimicking conditions in living cells, where much of lipid transport occurs at membrane contact sites^3, 12, 24–26, 39, 40^. Tethering can be achieved by protein domains that flank the lipid transport module or by an additional protein added to the mixture. Protein-mediated liposome tethering, however, leads to aggregation and does not allow controlling the distance between bilayer. Given the limitation of this system, we explored the possibility of using DNA nanotechnology (specifically DNA origami) to develop an aggregation-independent lipid-transport assay, in which the distance between liposomes could be tightly controlled. To this aim, we capitalized on a recently reported methodology to generate small liposomes within DNA rings^33, 34^ and further developed this method to obtain pairs of such liposomes with a well-defined distance between them.

We first constructed a DNA-based scaffold capable of holding two DNA rings at a precisely defined distance. Briefly, we designed DNA rings for templating size-defined liposomes as described previously^33, 34^. A ring (inner diameter ≈ 49 nm) consisted of a bundle of eight curved DNA helices with two of the inner helices lined by a total of 32 single-stranded (ss) oligonucleotide handles, which can hybridize to lipidated oligonucleotide anti-handles to nucleate the formation of liposomes (Fig. 1a and Supplementary Fig. 1). In our new design, each ring was connected to a six-helix-bundle DNA rod (130, 46, or 23 nm long) equipped with a sticky free end (Fig. 1b). These DNA nanostructures were allowed to fold by thermal annealing, purified by rate-zonal centrifugation^41^, and examined by negative-staining transmission electron microscopy (TEM), which confirmed the formation of rings connected to a rod (Fig. 1a). Next, DNA rings carrying rods with complementary sticky ends (DNA-A and DNA-B) were incubated for 90 min at room temperature (RT) to form dumbbell-like dimers, where two rings are connected by a pillar, as confirmed by TEM and agarose gel electrophoresis (Fig. 1b and Supplementary Fig. 2).

**Figure 1.**
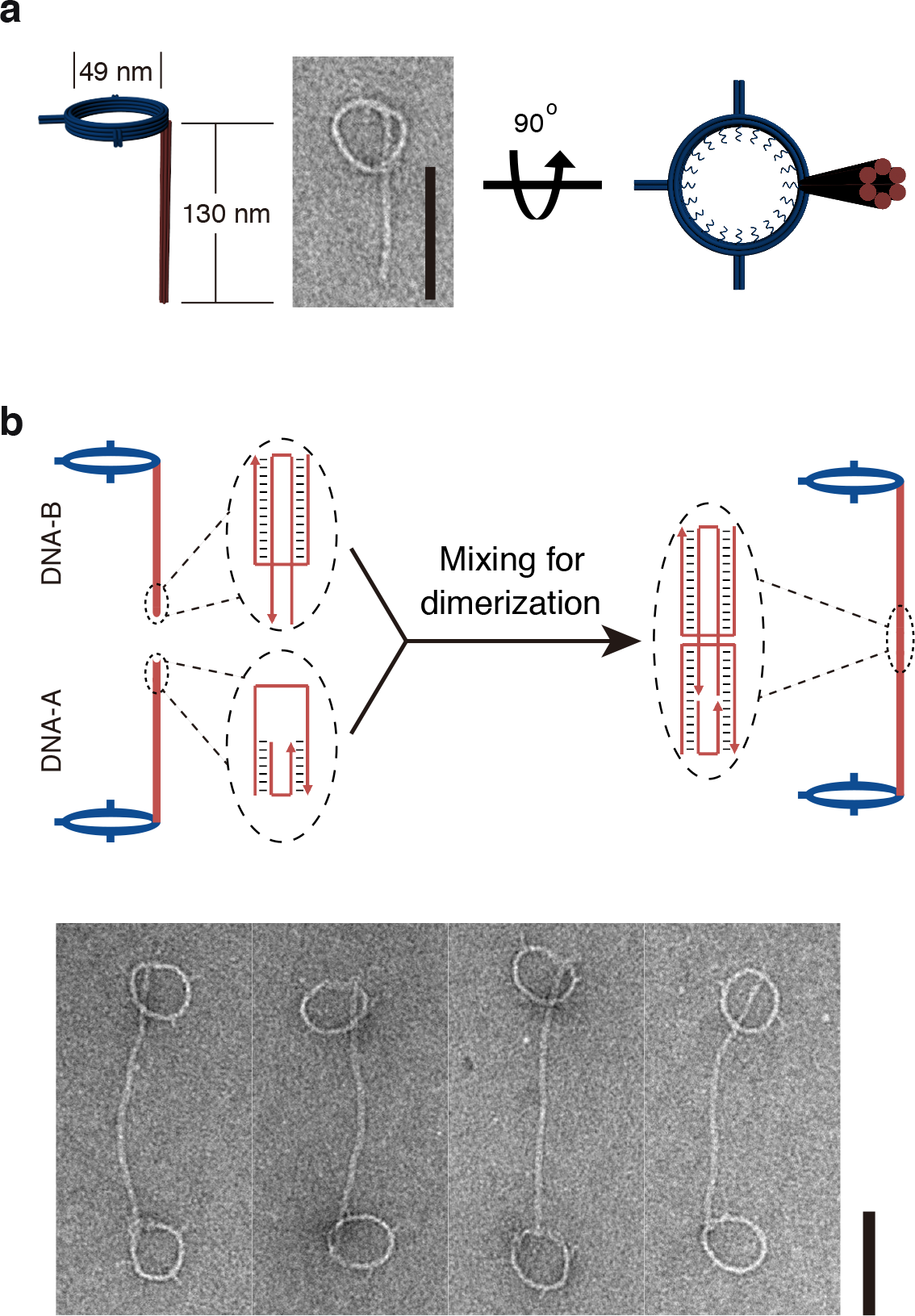
DNA nanostructures designed for this study. **a**, Cartoon models and a negative-staining TEM image of monomeric DNA nanostructure comprising a ring connected to a rod. **b**, Schematics of DNA nanostructure dimerization. Diagrams in the dashed ovals illustrate the dimerization mediated by DNA rods with complementary sticky ends. Negative-stain TEM images of DNA-origami dimers (two rings joint by a pillar) are shown at the bottom. Scale bars: 100 nm.

To produce dimeric DNA nanostructures harboring liposomes of different lipid compositions, monomeric DNA rings were labeled with phospholipids through inner handle/anti-handle hybridization, incubated with additional lipids and detergent, and dialyzed against detergent-free buffer solutions to allow liposome formation within the DNA rings (Fig. 2a)^33, 34^. Two sets of DNA ring-templated liposomes with different lipid compositions were generated and purified by iodixanol-gradient centrifugation. One set, termed “donor” liposomes (formed on DNA rings carrying DNA-A-labeled rods), contained DOPC, DGS-NTA(Ni), and a pair of fluorescence resonance energy transfer (FRET) dye-labeled lipids: NBD-PE and rhodamine-PE. The other set, termed “acceptor” liposomes (formed on DNA rings carrying DNA-B-labeled rods), contained DOPC, POPS, and PI(4,5)P_2_ (see Online Methods for lipid acronyms and molar ratios). DNA ring-templated donor and acceptor liposomes were then mixed to allow dimerization via DNA-A/DNA-B hybridization. Negative-staining TEM showed liposome-containing DNA-origami dimers (Fig. 2b and Supplementary Fig. 3) with center-to-center distance of the two liposomes consistent with the theoretical length of the DNA pillar. For example, using DNA rings with 130-nm rods, we obtained a liposome center-to-center distance of 252 ± 23 nm (Fig. 2d), confirming that the dimerized DNA pillars (~261 nm) was rigid enough to keep the two liposomes at the predicted distance. As the DNA ring-templated liposomes were homogenous in size and on average ~28 nm in diameter^34^, the closest distance between their membranes was on average ~224 nm. Similarly, when the donor and accepter liposomes were separated by 93-nm and 45-nm DNA pillars (Fig. 2c), TEM revealed liposome center-to-center distances of 95 ± 20 nm and 54 ± 17 nm, respectively. These values were again in good agreement with the design (Fig. 2d) and translated to membrane separations of circa 67 nm and 26 nm, respectively. We next used these donor/acceptor liposome pairs with controlled bilayer distances for lipid-transfer assays.

**Figure 2.**
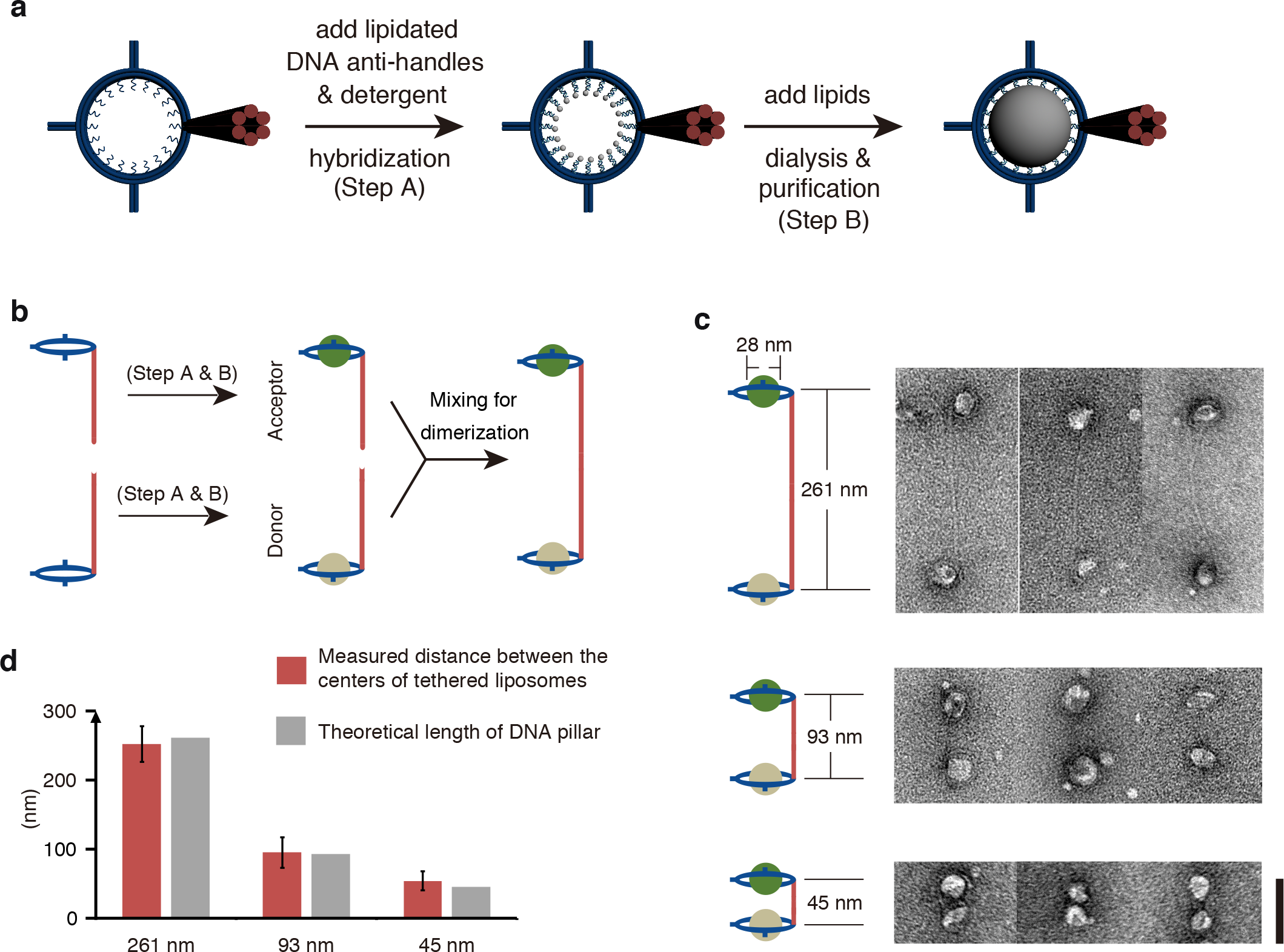
DNA-directed liposome formation and assembly. **a**, Schematics of DNA ring-templated liposome formation. **b**, Schematics of the donor and acceptor liposome heterodimer assembly. **c**, Cartoon models and TEM images of DNA-organized liposome dimers separated by DNA pillars with different lengths (261, 93, and 45 nm). **d**, Center-to-center distance of DNA-tethered liposomes measured from negative-stain TEM images compared with theoretical lengths of DNA pillars. Error bars represent SD (N=50, 50, and 25). Scale bars: 100 nm.

### Validation of DNA-templated liposomes for lipid transfer studies

As a premise to the use of dimeric DNA nanostructures to investigate the impact of bilayer distance on lipid transport by the SMP domain of E-Syt1, we first assessed whether presence of DNA rings affected the property of the SMP domain to transfer lipids between them. In this system, a small portion of the liposome surface is in contact with the negatively charged DNA nanostructure, and this could impact the properties of the bilayer. Furthermore, DNA-liposomes nanostructures must be maintained in a buffer that differs from those typically used for lipid-transport assays as it contains 10 mM Mg^2+^ (rather than 0 mM Mg^2+^) and 400 mM KCl (rather than 100 mM NaCl/KCl). Therefore, effects of these parameters need to be experimentally examined.

To address the potential impact of such parameters in our assay, lipid transfer with either DNA-free liposomes or DNA-templated liposomes was compared using the entire cytosolic region of E-Syt1. E-Syt1 with an N-terminal His-tag (E-Syt1_cyto_) to mediate binding to the DGS-NTA(Ni) containing liposomes was added to a mixture of DNA-free or DNA-templated donor and acceptor liposomes (Fig. 3a). Next, Ca^2+^ was also added as it was required to enable liposome tethering by the C2 domains of E-Syt1 and to relieve their inhibitory function on lipid transport^24–26^. Thus, in this system, E-Syt1_cyto_ acted both as the tether (His-tag and C2 domains) and as the lipid transport protein (SMP domain). As expected, addition of E-Syt1_cyto_ resulted in an increase in turbidity of the liposome mixture, which was revealed by optical density reading at 405 nm (turbidity) (Fig. 3b), reflecting E-Syt1_cyto_ - dependent aggregation of liposomes into large particles. Importantly, lipid transport was also observed under these conditions, as the fluorescence of NBD, which was quenched by rhodamine in the donor liposomes, increased due to dequenching, revealing dilution of NBD-PE and rhodamine-PE as they were transferred to acceptor liposomes. Transfer occurred with a slower kinetic with DNA-templated liposomes than with DNA-free liposomes, possibly reflecting, at least in part a delay in tethering (Fig. 3c, compare with Fig. 3b). In spite of the slower transfer, however, these results validate the use of our experimental system for investigations of lipid transport between bilayers.

**Figure 3.**
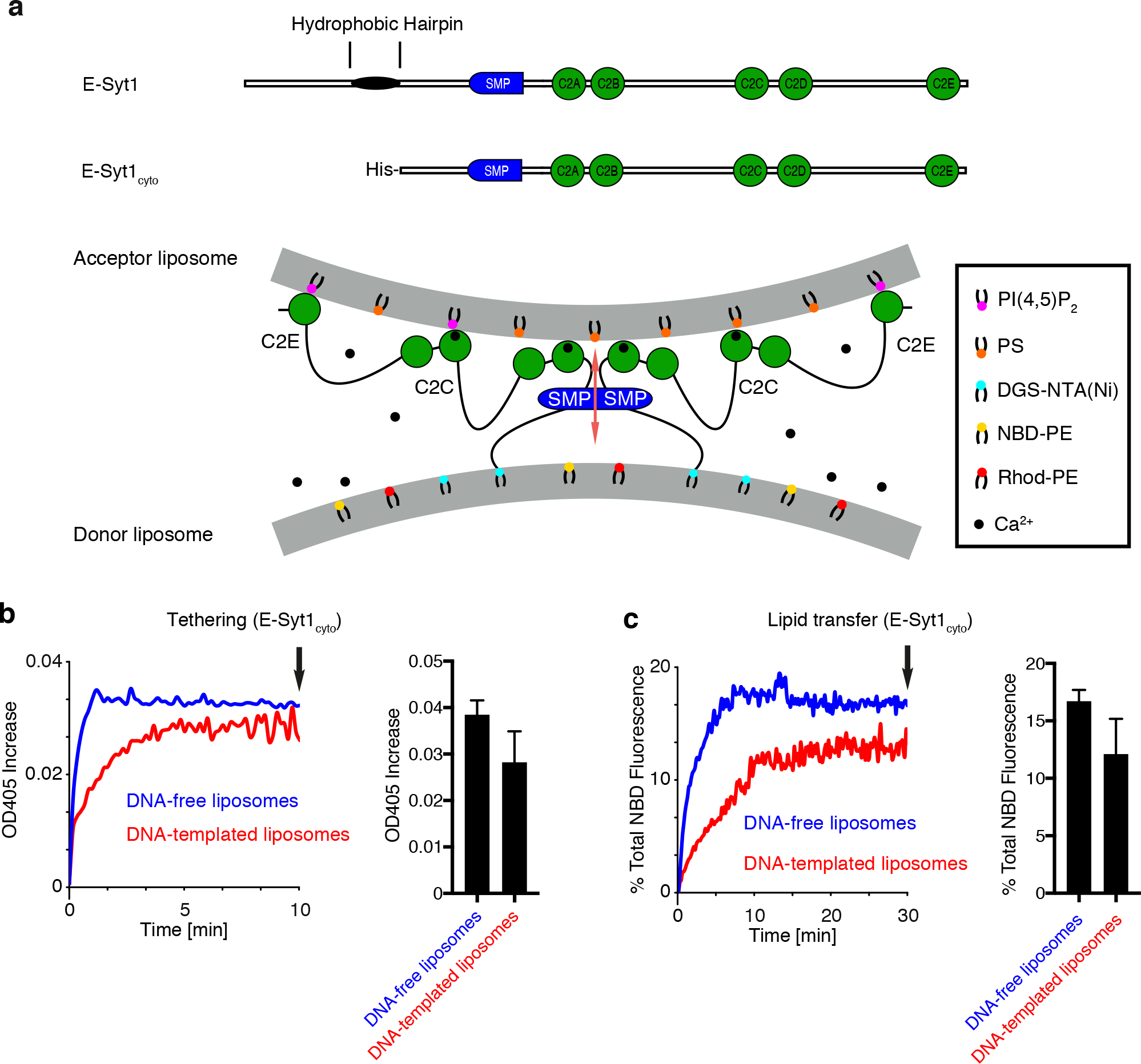
E-Syt1_cyto_ clusters and mediates lipid transfer between DNA origami templated liposomes. **a**, Domain structures of E-Syt1 and E-Syt1_cyto_ (top) and schematic representation of E-Syt1_cyto_-mediated liposome tethering and lipid transfer in the presence of Ca^2+^ (bottom). **b**, (Left) Time-courses of tethering of donor and acceptor liposomes in the presence of E-Syt1_cyto_ and Ca^2+^, as assessed by the increase in turbidity (OD at 405 nm). (Right) Bar graphs showing OD_405_ levels after incubation for 10 min (arrow in the left panel). **c**, (Left) Time-courses of lipid transfer between donor and acceptor liposomes in the presence of E-Syt1_cyto_ and 100 *μ*M Ca^2+^ at RT as assessed by the dequenching of NBD-PE fluorescence. (Right) Bar graphs showing quantification of NBD fluorescence at the end of the incubation (arrow in the left panel). Bar graphs in (b) and (c) show mean and SD of three independent experiments.

### DNA-organized liposome dimers as a model system to assess the distance at which lipid transport between bilayers can occur

To assess whether SMP domains can transport lipids by shuttling between the two liposomes within DNA-organized liposome dimers, we used constructs comprising the SMP domain of E-Syt1 and upstream “linker” sequences of different length. These constructs, which were preceded by a His-Tag, were devoid of the entire downstream C2 domain-containing region of the protein to prevent E-Syt1-dependent liposome aggregation, thus limiting our interrogation to lipid transport within individual DNA-templated liposome dimers (Fig. 4a). In one construct, SMP-native (SMP-N), the upstream linker region consisted of the 47 amino acid (a.a.) sequence (predicted to be unfolded) that connects the SMP domain to the hydrophobic hairpin region in wild-type E-Syt1, plus 35 a.a. derived from the vector. In another construct, SMP-short (SMP-S), the linker region comprised the SMP domain flanked at its N-terminus only by the 16 a.a. derived from the vector. In a third construct, SMP-long (SMP-L), the native a.a. sequence was replaced by a concatemer of 4 such sequences, resulting in a 214 a.a. linker. We used a concatemer of the native sequence for this longer construct to minimize the chance that this sequence could negatively impact the function of the SMP domain.

**Figure 4.**
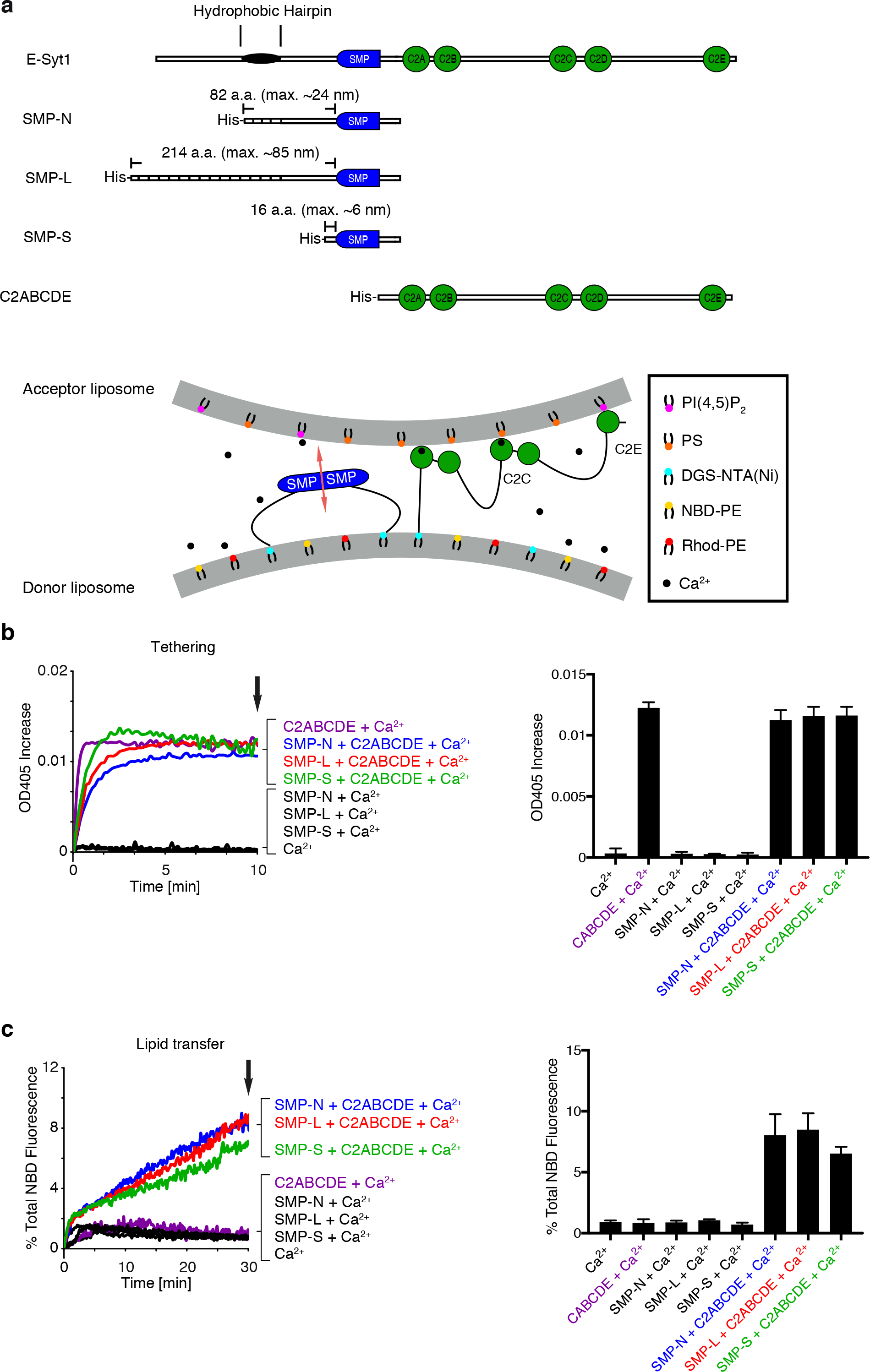
The SMP domain of E-Syt1 alone does not cluster liposomes but transfers lipids between protein-tethered liposomes. **a**, Domain structures of E-Syt1 constructs (top) and schematic representation of SMP-mediated lipid transfer between liposomes tethered by the C2ABCDE of E-Syt1 in the presence of Ca^2+^ (bottom). **b**, (Left) Time courses of liposomes aggregation (or the lack thereof), due to tethering of donor and acceptor liposomes in the presence of various E-Syt1 constructs and 100 *μ*M Ca^2+^ at RT, as assessed by the increase in turbidity (OD 405 nm). (Right) Bar graphs showing OD_405_ levels after incubation for 10 min (arrow in the left panel). **c**, (Left) Time-courses of lipid transfer between donor and acceptor liposomes in the presence of E-Syt1 constructs and 100 *μ*M Ca^2+^ at RT as assessed by the dequenching of NBD-PE fluorescence. (Right) Bar graphs showing quantification of NBD fluorescence at the end of the incubation (arrow in the left panel). Bar graphs in (b) and (c) show mean and SD of three independent experiments.

Prior to performing lipid transfer studies using these constructs, it was important to confirm that all three constructs did not tether liposomes, thus ensuring that liposome pairs were kept in close proximity only with the help of DNA nanostructures, and not by liposome aggregation. A turbidity assay confirmed that this was indeed the case, whereas the same assay performed with the additional C2ABCDE fragment of E-Syt1 resulted in major turbidity increases, reflecting robust aggregation (Fig. 4b). As expected, SMP domain-dependent lipid transfer, which was detected by fluorescence increase, occurred only when liposomes were tethered, i.e. in the presence of the C2ABCDE fragment (Fig. 4c). Further, upstream linkers of different length led to similar lipid-transfer activities, confirming that linker constructs had negligible influence on the function of SMP domain.

We next examined SMP-N-mediated lipid transfer using DNA-templated liposome dimers with a pillar length of 45 nm (dubbed 45-nm DNA-liposome in Fig. 5). In such dimers, the average distance between the bilayers of the two liposomes is estimated to be ~26 nm, which corresponds to a typical ER-PM contact distance (10–30 nm)^22^. The SMP-domain constructs were first added to the monomeric DNA templated donor liposomes to allow its binding to DGS-NTA(Ni) present in their bilayers. Subsequently, these liposomes were mixed with the monomeric DNA-templated acceptor liposomes for 90 min to allow dimer formation and lipid transport (Fig. 5a). A robust increase in fluorescence was observed, signaling the occurrence of lipid transfer (Fig. 5b). Consistent with the reported property of the E-Syts to harbor and/or transfer glycerolipids between membrane bilayers without major selectivity for specific head groups^10^, a similar increase in fluorescence was observed when NBD-PE in the donor liposomes was replaced by NBD-PS (Fig. 5c).

**Figure 5.**
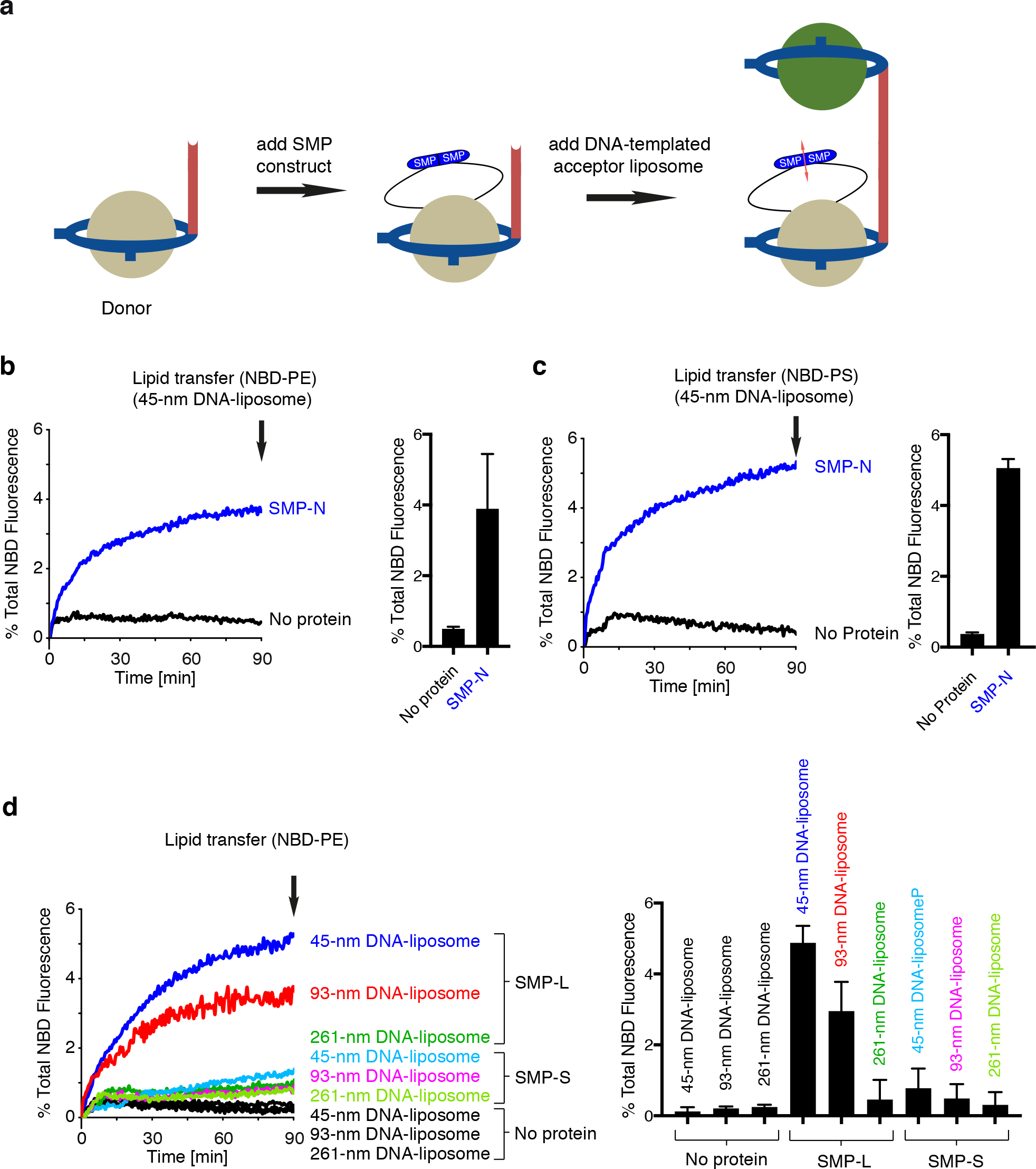
SMP mediates lipid transfer between the two liposomes within DNA-organized liposomes dimers. **a**, Schematic representation of the experimental protocol. **b**, Lipid transfer between donor and acceptor liposomes tethered by 45-nm DNA pillars in the absence or presence of the SMP-N at RT, as assessed by the dequenching of NBD-PE fluorescence. Time courses are on the left and bar graphs showing the NBD fluorescence at the end of the incubation (arrows in the left panel) are on the right. **c**, Same as in (b), but using donor liposomes in which NBD-PE was replaced by NBD-PS. **d**, Same as in (b), but liposomes were tethered by DNA pillars with greater lengths (93 and 261 nm) and transport was mediated by SMP constructs with two different polypeptide linkers (SMP-L, 214 a.a. and SMP-S, 16 a.a.). Mean and SD are derived from three independent experiments.

Next we tested SMP-S and SMP-L with dimeric DNA nanostructures comprising 45-, 93- and 261-nm DNA pillars (and thus bilayer distances of ~ 26, 67 and 224 nm). Similar to SMP-N, SMP-L, with a linker predicted to span 85 nm when fully stretched (Fig. 4a), showed robust lipid transfer between liposomes separated by the 45-nm pillars (Fig. 5d). More importantly, lipid transfer activity of SMP-L was observed when liposomes were separated by the 93-nm pillars (bilayer distance approximately 67 nm), but not by the 261-nm pillars (bilayer distance approximately 224 nm) (Fig. 5d). In contrast, shortening the linker region of the SMP domain to a maximum length of ~ 6 nm (SMP-S, Fig. 4a) abolished the lipid transfer activity on the 45-nm DNA-liposome dimers (~26 nm distance between bilayers) (Fig. 5d). This defect was not due to an impairment of the SMP domain per se, as SMP-S was functional on liposomes tethered by the C2ABCDE construct (Fig. 4).

These findings further indicate that exchange of lipids occurs within DNA-organized liposome dimers and validate the suitability of our nano-engineered system to study distance-dependent lipid transport between bilayers. Importantly, our data support the hypothesis that the SMP domain of E-Syts can act as a shuttle to move lipid across apposing bilayers (Fig. 6).

**Figure 6.**
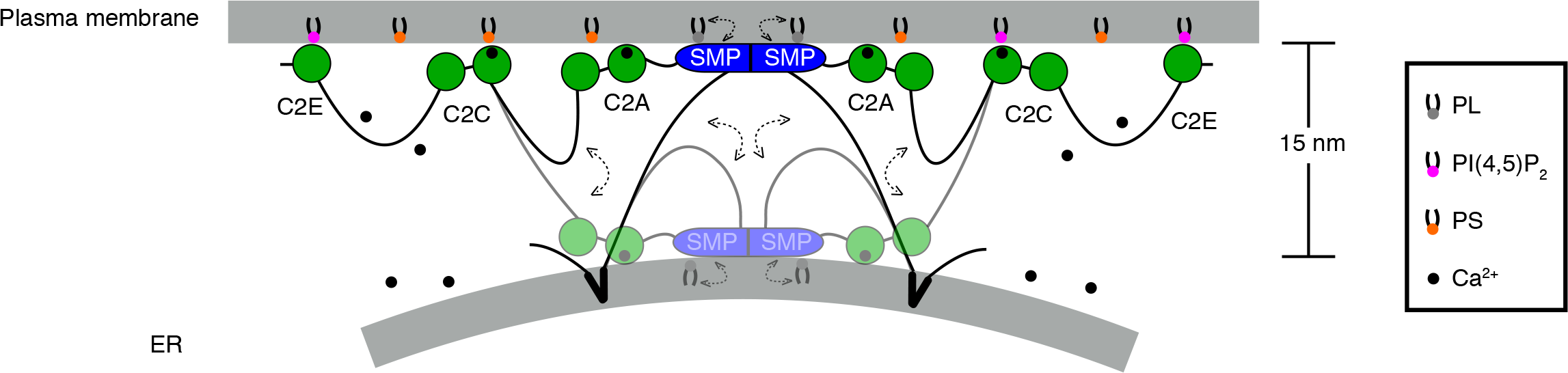
Model for E-Syt1-mediated lipid transfer. The interaction of E-Syt1 with the plasma membrane mediated by its C2C and C2E domain allows E-Syt1 to function as a tether at ER-plasma membrane contact sites. Both these interactions are triggered by elevation of cytosolic Ca^2+^.^20, 22–25, 35^ The SMP domain shuttles between the two membranes to transfer lipids. The Ca^2+^-binding C2A domain, which is tightly conjoined with the C2B domain, is theorized to move together with SMP domain. Ca^2+^ ions are shown as small black circles. Double-headed dashed arrows indicate the movement of SMP domain together with the C2A-C2B domain module.

## Discussion

DNA nanotechnology has allowed for the construction of precisely engineered lipid bilayer structures that are amenable to biochemical assays (e.g. FRET-based analyses), thus providing a unique opportunity to study membrane dynamics quantitatively and systematically^32^. In this work, we joined two DNA ring-templated liposomes with distinct lipid compositions via a rigid DNA pillar. Unlike liposomes tethered by proteins, which cluster into tight aggregates, here the two closely apposed membrane bilayers can be maintained at customized distances with nanoscale precision by programmable DNA nanostructures. Thus, the DNA-origami-directed membrane engineering technique offers a solution to generate membrane appositions *in vitro* with controllable membrane proximity.

Here we have used DNA origami-organized liposomes to assess whether the SMP domain of E-Syt1 can transfer lipids at a distance that exceeds its length. In our experimental system, membrane-tethering via C2 domains was replaced by membrane tethering via DNA nanostructures, allowing to control liposome separation distance in a pre-defined, non-aggregating way. An important issue of these studies was the demonstration that the FRET signal changes observed in our assay reflected lipid transfer between pairs of liposomes within dimeric DNA nanostructures. This was found to be the case as only minimal FRET signal changes occurred with untethered liposomes by SMP domain. Please note that the average distance of untethered liposomes in suspension under the conditions of our assay - liposome size and lipid concentration (0.1-0.5 mM) - is estimated to be around 403-709 nm^42^.

We show that upon anchoring to the donor liposomes via unstructured linker regions of different length, the SMP domain of E-Syt1 can transfer lipids with the dimeric structures at distances that greatly exceed the length of dimeric SMP domain (~ 9 nm) but are not longer than the length of those linker regions. The shuttle model is consistent with the hydrophilic nature of the two tips of the SMP domain and with the lack of evidence for the property of the SMP domain to form tetramers or longer adducts to bind and bridge the two bilayers^10^. The polypeptide sequence that links the SMP domain of E-Syt1 to its intramembrane hydrophobic hairpin (47 a.a.) is predicted to be unfolded. With the maximal theoretical length approaching 19 nm, this linker is long enough to allow the SMP domain to shuttle lipids over the inter-membrane distance typically found between the ER and the plasma membrane (~15 nm) in cells expressing E-Syt1 under high cytosolic Ca^2+^, where recruitment of E-Syt1 to the plasma membrane and its lipid transfer functions are stimulated^20–25^.

A shuttling of the SMP domain of E-Syt1 over an approximately 15 nm distance, however, raises the conundrum of how such shuttling can be compatible with the presence at the C-terminal side of the SMP domain of a closely adjacent C2 domain (C2A domain) with membrane binding properties^18, 24^. The C2A domains of the E-Syts bind membranes in a Ca^2+^-dependent way^18, 24, 43^and were proposed to bind the plasma membrane^44^. However, the length of the unstructured linker that separates it from the SMP domain is too short (~ 8 nm when fully stretched) to span the ER-plasma membrane distance when the SMP domain is in close proximity of the ER during its shuttling. Thus, we propose that in living cells the tandem arranged C2A-C2B domains^10, 44^ move together with the SMP domain between the two membranes during lipid transport (Fig. 6). In support of this possibility, membrane binding by the C2A domain is not selective for negatively charged bilayers, thus making plausible its binding to both bilayers^18, 24^. Its proximity to the SMP domain may help orient the SMP domain at the membrane interface for lipid extraction/delivery. Clearly, elucidating the dynamics of E-Syt-mediated lipid transfer warrants further investigation in the future. Outstanding questions include how and at what pace the SMP domain extracts lipids from membrane bilayers and delivers them to the target bilayers. A combination of our platform with single-molecule experiments could shed light on these questions.

Our current study with the SMP domain of E-Syt1 validates the concept of using DNA-origami-guided membrane structures to modulate and study lipid transport. In theory, this system can be adapted to study other lipid transport proteins as well as enzymatic reactions thought to involve “in trans” interactions of two membranes^45, 46^. Modification of the nanostructures used here will enable the generation of different membrane curvature and/or topology. Finally, it may be possible to use these nanostructures to aid the structural analysis of tethering proteins by cryo-EM.

## Online Methods

### Chemicals

Chemicals were from the following sources: Isopropyl-β-D-thiogalactoside (IPTG) (AmericanBio), His60 Ni Superflow Resin (Takara), Proteinase K (Sigma), tris (2-carboxyethyl) phosphine (TCEP) (Thermo), Iodixanol solution (60%, w/v) (Cosmo Bio USA), Octyl β-D-glucopyranoside (OG) (EMD Millipore), n-dodecyl-β-D-maltopyranoside (DDM) (Avanti Polar Lipids). All DNA oligonucleotides were purchased from Integrated DNA Technologies. All lipids were obtained from Avanti Polar Lipids: 1,2-dioleoyl-*sn*-glycero-3-phosphocholine (DOPC); 1-palmitoyl-2-oleoyl-*sn*-glycero-3-phospho-L-serine (POPS); L-α-phosphatidylinositol-4,5-bisphosphate [PI(4,5)P_2_]; 1,2-dioleoyl-sn-glycero-3-phosphoethanolamine-N-(7-nitro-2-1,3-benzoxadiazol-4-yl) (NBD-PE); 1,2-dioleoyl-sn-glycero-3-phospho-L-serine-N-(7-nitro-2-1,3-benzoxadiazol-4-yl) (NBD-PS); 1,2-dioleoyl-sn-glycero-3-phosphoethanolamine-N-(lissamine rhodamine B sulfonyl) (Rhod-PE); 1,2-dioleoyl-sn-glycero-3-[(N-(5-amino-1-carboxypentyl) iminodiacetic acid) succinyl] [DGS-NTA(Ni)]; 1,2-dioleoyl-sn-glycero-3-phosphoethanolamine-N-[4-(p-maleimidophenyl)butyramide] (MPB-PE).

### Plasmids

The plasmids encoding E-Syt1_cyto_ and C2ABCDE were previously described^24^. The region coding for the SMP domain of E-Syt1 plus its upstream sequence up to the hydrophobic hairpin (a.a.93-327 of E-Syt1, referred to here as SMP-N) was cloned using NotI and XhoI sites into the pET-28a vector (Novagen). SMP-S (a.a.135-327 of E-Syt1) was cloned using NdeI and NotI sites into the pET-28a vector. SMP-L including 3x a.a.93-136 and 93-327 was cloned into the pET-28a vector using BamHI, SacI, SalI, NotI and XhoI sites, respectively.

### Protein Expression and Purification

Soluble fragments of human E-Syt1 were expressed in Expi293 cells or BL21 (DE3) RIL Codon Plus (Agilent) *E. coli* cells and purified as described previously^24^. Briefly, cells were harvested and lysed in buffer A [25 mM Tris-HCl, pH 8.0, 300 mM NaCl, 10 mM imidazole, 1x complete EDTA-free protease inhibitor cocktail (Roche), 0.5 mM TCEP] by three freeze–thawing cycles using liquid nitrogen (for Expi293 cells) or by sonication (for bacteria). The suspension was clarified by centrifugation at 17,000 x g for 30 min at 4 °C and the protein was purified from the supernatant by a Ni-NTA column. After elution, the protein was further purified by gel filtration. Fractions containing E-Syt1 fragments were pooled and concentrated.

### Liposome Preparation

DNA-free donor liposomes were composed as follows: 87:1.5:1.5:10 mole percent of DOPC: NBD-PE: Rhod-PE: DGS-NTA(Ni) or 87:1.5:1.5:10 mole percent of DOPC: NBD-PS: Rhod-PE: DGS-NTA(Ni). DNA-free acceptor liposomes were composed as follows: 85:10:5 mole percent DOPC: POPS: PI(4,5)P_2_. Liposome preparation was performed as previously described^24^. Briefly, lipid mixtures were dried onto a film. Lipid films were then hydrated with buffer A. DNA-free liposomes were formed by ten freeze-thaw cycles in liquid N_2_ and 37 °C water bath and extrusion through polycarbonate filters with a pore size of 50 nm (Avanti Polar Lipids).

### DNA Origami Design and Preparation

Monomeric DNA-origami cages were designed using caDNAno (cadnano.org). Inner and outer handle sequences (21-nt) were generated by NUPACK (nupack.org) and manually added to the 3'-ends of the appropriate staple strands. DNA scaffold strands (8064-nt) were produced using *E. coli* and M13-derived bacteriophages^28, 30^. The DNA cage monomers were assembled from a scaffold strand (20 nM) and a pool of staple strands (140 nM each) in buffer B (5 mM Tris-HCl, pH 8.0, 12 mM of MgCl_2_) using a 36-hour thermal annealing program. Correctly assembled DNA cages were purified via rate-zonal centrifugation in glycerol gradients as described previously^34^.

### Preparation of lipidated DNA anti-handles

The lipid-DNA conjugate was prepared as previously described with slight changes^34^. Briefly, 1 mM 5’-thiol labeled DNA oligonucleotides were treated with 20 mM TCEP in buffer C (25 mM Hepes, pH 7.4, 140 mM KCl) for 30 min. 900 μM pre-treated thiol-modified DNA oligonucleotides were reacted with 3.3 mM pre-dried MPB-PE and 6.7 mM pre-dried donor lipid mixtures [77% DOPC, 20% DGS-NTA(Ni), 1.5% Rhod-PE, 1.5% NBD-PE] or pre-dried acceptor lipid mixtures [85% DOPC, 10% POPS, 5% PI(4,5)P_2_] in buffer C containing 2% OG at RT for 30 min. This reaction mixture was then dialyzed in 7 Kd molecular weight cut-off (MWCO) cassette against buffer C overnight in order to incorporate lipidated DNA molecules into liposomes. The conjugation products were separated from unconjugated DNA via isopycnic centrifugation in iodixanol gradients and later analyzed by SDS/PAGE. Fractions containing DNA-lipid conjugates were combined.

### Preparation of DNA Origami-Organized Liposomes

DNA origami-organized liposomes were prepared as previously described with slight changes^34^. Briefly, 20 nM assembled DNA nanostructures were first labeled with 1280 nM lipidated anti-handles buffer D (25 mM Hepes, pH 7.4, 400 mM KCl, 10 mM MgCl_2_) containing 1% OG at 37°C for 1h. To form DNA origami-organized liposomes, 0.6 mM rehydrated donor or acceptor lipid mixtures were added to 10 nM lipid-labeled DNA nanostructures. The solution was diluted to 500 μL in buffer D with 1% OG, gently shaken for 30 minutes at RT, put into a 7 kD MWCO dialysis cassette, and dialyzed against buffer D overnight. The dialyzed solutions were subjected to centrifugation in iodixanol gradients. A quasi-linear gradient containing 2.4 mL of 6%−26% (w/v) iodixanol was loaded on top of 2 mL dialyzed sample containing 30% iodixanol. Gradients were then centrifuged at 50,000 rpm for 5 h at 4°C. Fractions were analyzed by 1.5% agarose gel with 0.05% SDS and those containing DNA origami-organized liposomes were combined.

### Electron Microscopy

To prepare negatively stained DNA nanostructures or DNA origami-organized liposomes, a drop of sample (5 μL) was deposited on a glow discharged formvar/carbon coated copper grid and incubated for 1-3 min at RT. Fluid was then removed by blotting with a filter paper. The grid was immediately stained for 3 min with 2% (w/v) uranyl formate. Grids were examined using a JEOL JEM-1400 Plus microscope equipped with a LaB6 filament (acceleration voltage: 80 kV). Images were acquired by an Advanced Microscopy Technologies bottom-mount 4k×3k CCD camera.

### Analysis of DNA Nanostructures Dimerization Efficiency

5 μL of DNA nanostructures or DNA origami-organized liposomes were incubated with 0.2 μL 1 mM 3’-Cy3 labeled DNA oligonucleotides at 37 °C for 30 min. The samples were then analyzed by 1.5% agarose gel with 0.05% SDS and 10 mM MgCl_2_. Fluorescence of DNA bands was visualized using Typhoon FLA 9500 (GE Healthcare), and the fluorescence density were analyzed using ImageJ (NIH).

### Lipid Transfer Assays Using DNA-Free Liposomes

All *in vitro* lipid transfer assays with soluble proteins were performed as previously described^24^. Briefly, reactions were performed in 50 μL volumes with a final lipid concentration of 0.5 mM, with donor and acceptor liposomes added at a 1:1 ratio. Reactions were initiated by the addition of proteins to the liposome mixtures (protein: lipid ratio 1: 1000) in a 96-well plate (Corning). The fluorescence intensity of NBD was monitored with an excitation of 460 nm and emission of 538 nm every 10 sec over 30 min at RT by using SpectraMax M5 Microplate Reader (Molecular Devices). All data were corrected by setting the data point at 0 min to zero, and subtracting the baseline values obtained in the absence of proteins. The data were expressed as a percentage of the maximum fluorescence, determined after adding 10 μL of 2.5% DDM to the reactions after 30 min. All experiments were repeated 3 times and a representative trace is shown. Bars represent average fluorescence values +/− standard errors from the three-pooled readings.

### Lipid Transfer Assays Using DNA Origami-Organized Liposomes

Lipid transfer assays using DNA origami-organized liposomes were performed as for the lipid transfer assays using DNA-free liposomes, with the exception that1 μM SMP domain was preincubated with 100 μM DNA ring-templated donor liposomes for 5 min at RT. Reactions were initiated by the addition of 100 μM DNA ring-templated acceptor liposomes in a 96-well plate (Corning). The DDM detergent was added to stop the reactions after 90 min. All experiments were repeated 3 times and a representative trace is shown. Bars represent average fluorescence values +/− standard errors from the three-pooled readings.

### Liposome Tethering Assays

Liposome tethering assays with soluble proteins were performed as previously described^24^. Briefly, the reactions were initiated by the addition of proteins to the mixture of donor and acceptor liposomes (1:1 ratio) in a 96-well plate (Corning) using SpectraMax M5 Microplate Reader (Molecular Devices). The absorbance at 405 nm was measured to assess turbidity. Data were expressed as absolute absorbance values subtracted by the absorbance prior to protein addition. All experiments were repeated 3 times. Bars represent average 405 nm absorbance values from the three-pooled readings.

## Acknowledgments

We thank Yiying Cai for discussion. This work was supported in part by NIH grants NS036251 and DA018343, the HHMI and the Kavli Foundation to P.D.C., an NIH Director’s New Innovator Award (GM114830) and a Yale University faculty startup fund to C.L., and by a Human Frontier Science Program Long-term Fellowship to X.B..

## Author Contribution

All authors participated in the project initiation, data analysis and interpretation. X.B. designed and performed liposome tethering and lipid transfer assays, and prepared DNA origami-organized liposomes. Z.Z. designed the DNA origami, prepared DNA origami-organized liposomes and performed EM studies. X.B., Z.Z., P.D.C. and C.L. wrote the manuscript.

## Conflict of interest

The authors declare no conflict of interest.

## Figure Legends

**Supplementary Figure 1. Schematic illustration and caDNAno diagrams of DNA-origami nanostructures.** The full structure, which consists of an eight-helix-bundle (on square lattice) ring, 3 four-helix-bundle (on square lattice) side bars, and 2 six-helix-bundle (on honeycomb lattice) rods, is divided into individual parts due to various versions of pillars in different final constructs. **a,** Ring with side bars. **b,** Long rod (rod 1) and its self-dimerization into a 261-nm pillar. **c,** Middle rod (rod 2) and its self-dimerization into a 93-nm pillar. **d,** Short rod (a modified version of rod 2) and its self-dimerization into a 45-nm pillar.

**Supplementary Figure 2. Dimeric DNA-origami nanostructures. a**, Gallery of negative-stain TEM images showing dimerized DNA-origami nanostructures with 261-, 93- and 45-nm DNA pillars, respectively. Scale bars: 100 nm. **b**, The DNA-origami nanostructures were analyzed by SDS-agarose gel electrophoresis. The arrow and the triangle denote the monomeric and dimeric DNA structures, respectively. The dimer/monomer ratios, as revealed by the densitometry, are shown at the bottom.

**Supplementary Figure 3. DNA-organized liposome dimers. a**, Gallery of negative-stain TEM images showing liposome dimers separated by 261-, 93- and 45-nm DNA pillars, respectively. Scale bars: 100 nm. **b**, The DNA-organized liposome dimers were analyzed by SDS-agarose gel electrophoresis. The arrow and the triangle denote the monomeric and dimeric DNA structures, respectively. The dimer/monomer ratios, as revealed by densitometry, are shown at the bottom.

## References

1. Saheki, Y. & De Camilli, P. Endoplasmic Reticulum-Plasma Membrane Contact Sites. Annual review of biochemistry (2017).

2. Wu, H., Carvalho, P. & Voeltz, G.K. Here, there, and everywhere: The importance of ER membrane contact sites. Science 361(2018).

3. Wong, L.H., Gatta, A.T. & Levine, T.P. Lipid transfer proteins: the lipid commute via shuttles, bridges and tubes. Nat Rev Mol Cell Biol (2018).

4. Stefan, C.J., Manford, A.G. & Emr, S.D. ER-PM connections: sites of information transfer and inter-organelle communication. Curr Opin Cell Biol 25, 434–442 (2013).

5. AhYoung, A.P. et al. Conserved SMP domains of the ERMES complex bind phospholipids and mediate tether assembly. Proceedings of the National Academy of Sciences of the United States of America 112, E3179–3188 (2015).

6. Jeong, H., Park, J., Jun, Y. & Lee, C. Crystal structures of Mmm1 and Mdm12-Mmm1 reveal mechanistic insight into phospholipid trafficking at ER-mitochondria contact sites. Proceedings of the National Academy of Sciences of the United States of America 114, E9502–E9511 (2017).

7. Kawano, S. et al. Structure-function insights into direct lipid transfer between membranes by Mmm1-Mdm12 of ERMES. The Journal of cell biology 217, 959–974 (2018).

8. Kopec, K.O., Alva, V. & Lupas, A.N. Homology of SMP domains to the TULIP superfamily of lipid-binding proteins provides a structural basis for lipid exchange between ER and mitochondria. Bioinformatics 26, 1927–1931 (2010).

9. Lee, I. & Hong, W. Diverse membrane-associated proteins contain a novel SMP domain. FASEB journal : official publication of the Federation of American Societies for Experimental Biology 20, 202–206 (2006).

10. Schauder, C.M. et al. Structure of a lipid-bound extended synaptotagmin indicates a role in lipid transfer. Nature 510, 552–555 (2014).

11. Wong, L.H. & Levine, T.P. Tubular lipid binding proteins (TULIPs) growing everywhere. Biochimica et biophysica acta. Molecular cell research 1864, 1439–1449 (2017).

12. Lees, J.A. et al. Lipid transport by TMEM24 at ER-plasma membrane contacts regulates pulsatile insulin secretion. Science 355(2017).

13. Alva, V. & Lupas, A.N. The TULIP superfamily of eukaryotic lipid-binding proteins as a mediator of lipid sensing and transport. Biochimica et biophysica acta 1861, 913–923 (2016).

14. Kornmann, B. et al. An ER-mitochondria tethering complex revealed by a synthetic biology screen. Science 325, 477–481 (2009).

15. Liu, L.K., Choudhary, V., Toulmay, A. & Prinz, W.A. An inducible ER-Golgi tether facilitates ceramide transport to alleviate lipotoxicity. The Journal of cell biology 216, 131–147 (2017).

16. Toulmay, A. & Prinz, W.A. A conserved membrane-binding domain targets proteins to organelle contact sites. Journal of cell science 125, 49–58 (2012).

17. Hirabayashi, Y. et al. ER-mitochondria tethering by PDZD8 regulates Ca(2+) dynamics in mammalian neurons. Science 358, 623–630 (2017).

18. Min, S.W., Chang, W.P. & Sudhof, T.C. E-Syts, a family of membranous Ca2+-sensor proteins with multiple C2 domains. Proceedings of the National Academy of Sciences of the United States of America 104, 3823–3828 (2007).

19. Manford, A.G., Stefan, C.J., Yuan, H.L., Macgurn, J.A. & Emr, S.D. ER-to-plasma membrane tethering proteins regulate cell signaling and ER morphology. Developmental cell 23, 1129–1140 (2012).

20. Giordano, F. et al. PI(4,5)P(2)-dependent and Ca(2+)-regulated ER-PM interactions mediated by the extended synaptotagmins. Cell 153, 1494–1509 (2013).

21. Chang, C.L. et al. Feedback regulation of receptor-induced Ca2+ signaling mediated by E-Syt1 and Nir2 at endoplasmic reticulum-plasma membrane junctions. Cell reports 5, 813–825 (2013).

22. Fernandez-Busnadiego, R., Saheki, Y. & De Camilli, P. Three-dimensional architecture of extended synaptotagmin-mediated endoplasmic reticulum-plasma membrane contact sites. Proceedings of the National Academy of Sciences of the United States of America 112, E2004–2013 (2015).

23. Idevall-Hagren, O., Lu, A., Xie, B. & De Camilli, P. Triggered Ca2+ influx is required for extended synaptotagmin 1-induced ER-plasma membrane tethering. The EMBO journal 34, 2291–2305 (2015).

24. Bian, X., Saheki, Y. & De Camilli, P. Ca(2+) releases E-Syt1 autoinhibition to couple ER-plasma membrane tethering with lipid transport. The EMBO journal 37, 219–234 (2018).

25. Saheki, Y. et al. Control of plasma membrane lipid homeostasis by the extended synaptotagmins. Nature cell biology 18, 504–515 (2016).

26. Yu, H. et al. Extended synaptotagmins are Ca2+-dependent lipid transfer proteins at membrane contact sites. Proceedings of the National Academy of Sciences of the United States of America 113, 4362–4367 (2016).

27. Orci, L. et al. From the Cover: STIM1-induced precortical and cortical subdomains of the endoplasmic reticulum. Proceedings of the National Academy of Sciences of the United States of America 106, 19358–19362 (2009).

28. Douglas, S.M. et al. Self-assembly of DNA into nanoscale three-dimensional shapes. Nature 459, 414–418 (2009).

29. Rothemund, P.W. Folding DNA to create nanoscale shapes and patterns. Nature 440, 297–302 (2006).

30. Dietz, H., Douglas, S.M. & Shih, W.M. Folding DNA into twisted and curved nanoscale shapes. Science 325, 725–730 (2009).

31. Grome, M.W., Zhang, Z., Pincet, F. & Lin, C. Vesicle Tubulation with Self-Assembling DNA Nanosprings. Angewandte Chemie 57, 5330–5334 (2018).

32. Xu, W. et al. A Programmable DNA Origami Platform to Organize SNAREs for Membrane Fusion. Journal of the American Chemical Society 138, 4439–4447 (2016).

33. Yang, Y. et al. Self-assembly of size-controlled liposomes on DNA nanotemplates. Nature chemistry 8, 476–483 (2016).

34. Zhang, Z., Yang, Y., Pincet, F., Llaguno, M.C. & Lin, C. Placing and shaping liposomes with reconfigurable DNA nanocages. Nature chemistry 9, 653–659 (2017).

35. Chan, Y.H., van Lengerich, B. & Boxer, S.G. Effects of linker sequences on vesicle fusion mediated by lipid-anchored DNA oligonucleotides. Proceedings of the National Academy of Sciences of the United States of America 106, 979–984 (2009).

36. Franquelim, H.G., Khmelinskaia, A., Sobczak, J.P., Dietz, H. & Schwille, P. Membrane sculpting by curved DNA origami scaffolds. Nature communications 9, 811 (2018).

37. Perrault, S.D. & Shih, W.M. Virus-inspired membrane encapsulation of DNA nanostructures to achieve in vivo stability. ACS nano 8, 5132–5140 (2014).

38. Beales, P.A. & Vanderlick, T.K. Specific binding of different vesicle populations by the hybridization of membrane-anchored DNA. The journal of physical chemistry. A 111, 12372–12380 (2007).

39. Kumar, N. et al. VPS13A and VPS13C are lipid transport proteins differentially localized at ER contact sites. The Journal of cell biology 217, 3625–3639 (2018).

40. Mesmin, B. et al. A four-step cycle driven by PI(4)P hydrolysis directs sterol/PI(4)P exchange by the ER-Golgi tether OSBP. Cell 155, 830–843 (2013).

41. Lin, C., Perrault, S.D., Kwak, M., Graf, F. & Shih, W.M. Purification of DNA-origami nanostructures by rate-zonal centrifugation. Nucleic acids research 41, e40 (2013).

42. Xu, W., Wang, J., Rothman, J.E. & Pincet, F. Accelerating SNARE-Mediated Membrane Fusion by DNA-Lipid Tethers. Angewandte Chemie 54, 14388–14392 (2015).

43. Ma, L. et al. Single-molecule force spectroscopy of protein-membrane interactions. eLife 6(2017).

44. Xu, J. et al. Structure and Ca(2)(+)-binding properties of the tandem C(2) domains of E-Syt2. Structure 22, 269–280 (2014).

45. Stefan, C.J. et al. Osh proteins regulate phosphoinositide metabolism at ER-plasma membrane contact sites. Cell 144, 389–401 (2011).

46. Eden, E.R., White, I.J., Tsapara, A. & Futter, C.E. Membrane contacts between endosomes and ER provide sites for PTP1B-epidermal growth factor receptor interaction. Nature cell biology 12, 267–272 (2010).

