## Supplementary figures and images for "A programmable DNA-origami platform for studying protein-mediated lipid transfer between bilayers"

### Supplementary Figure 1

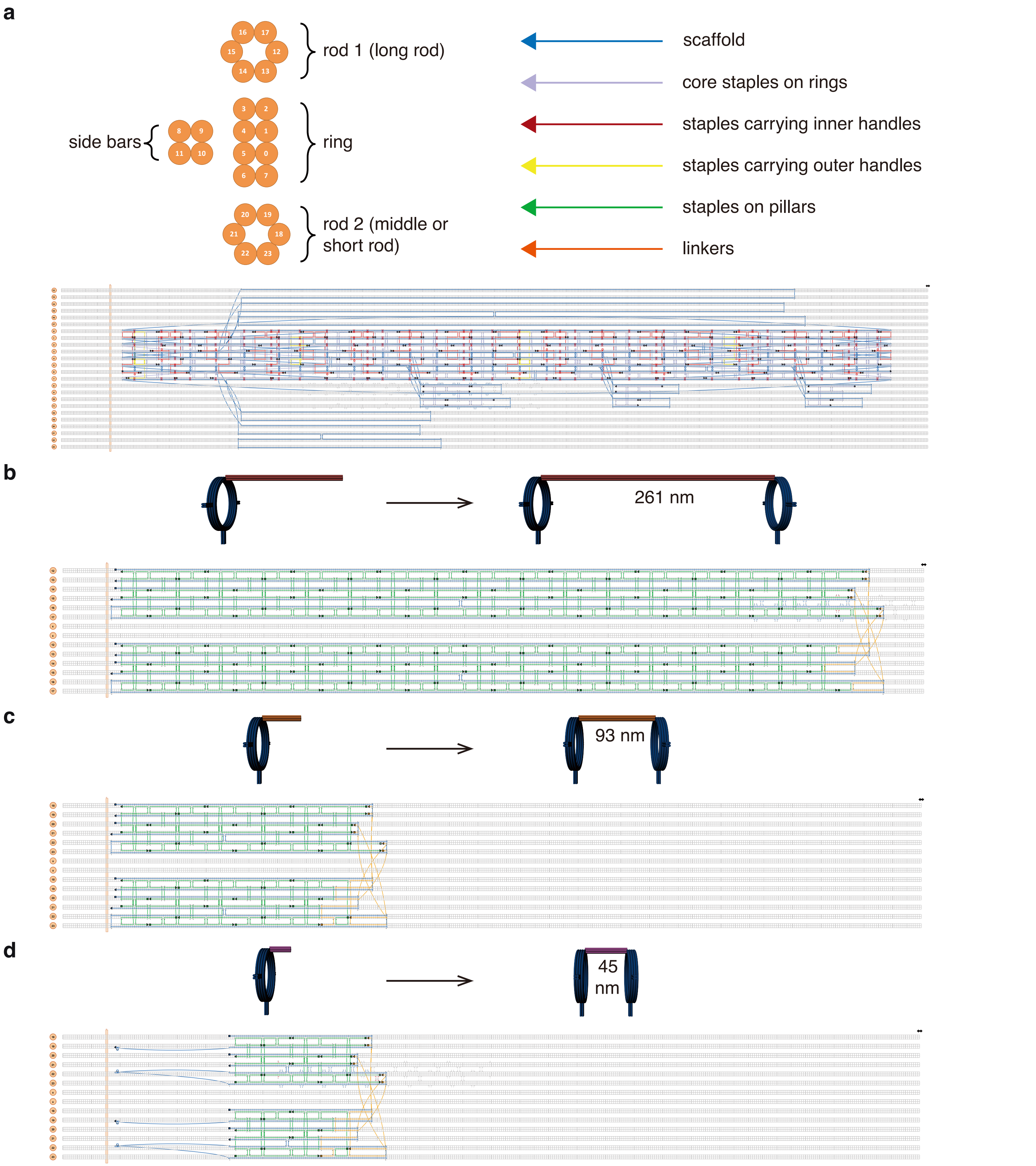


Supplementary Figure 1

### Supplementary Figure 2

**a**

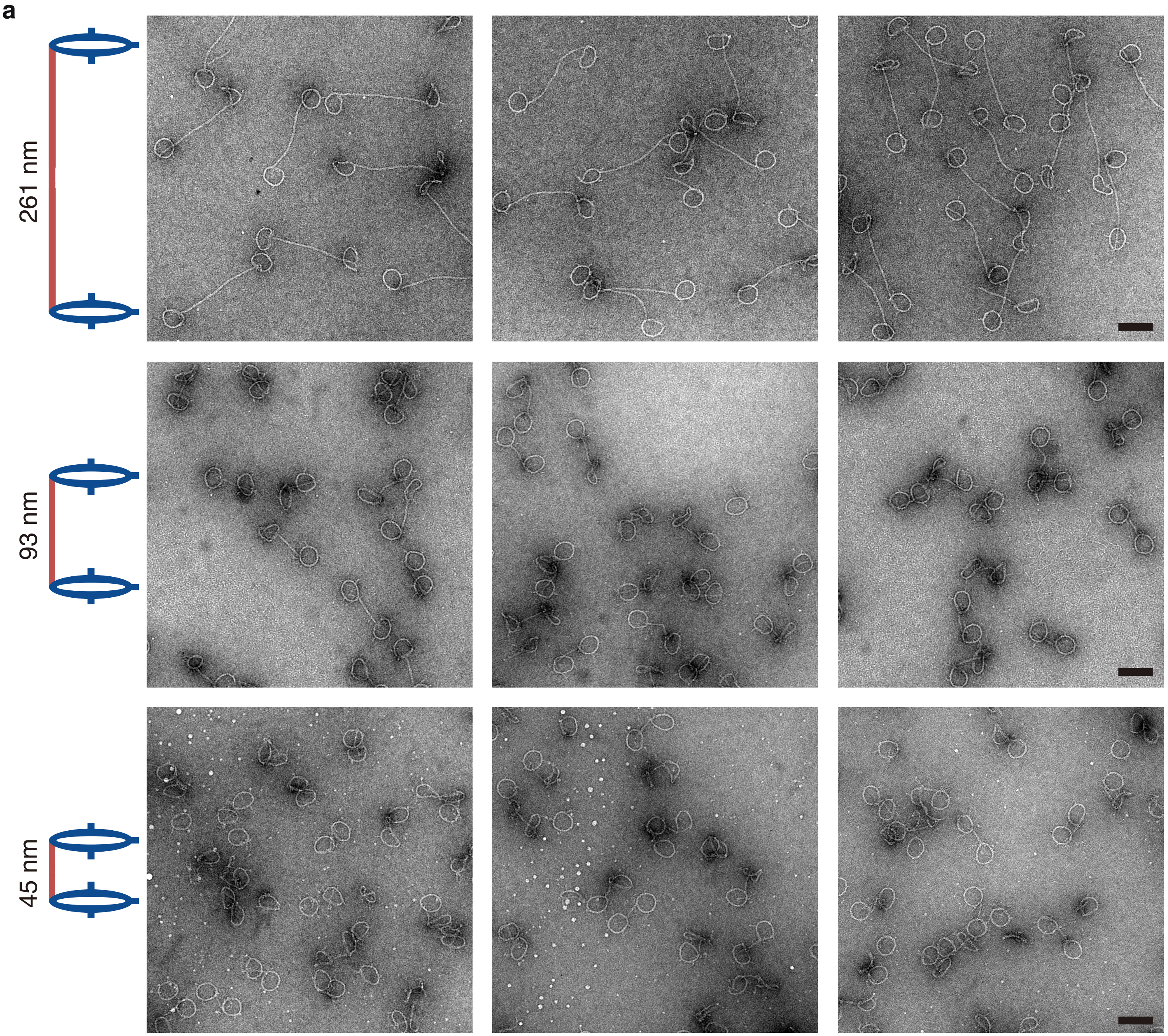


**b**

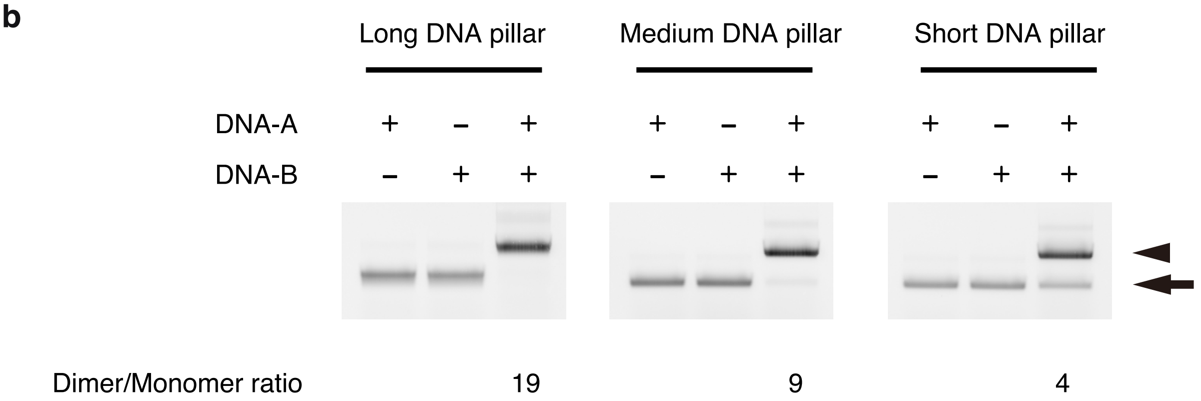


#### Supplementary Figure 2

### Supplementary Figure 3

**a**

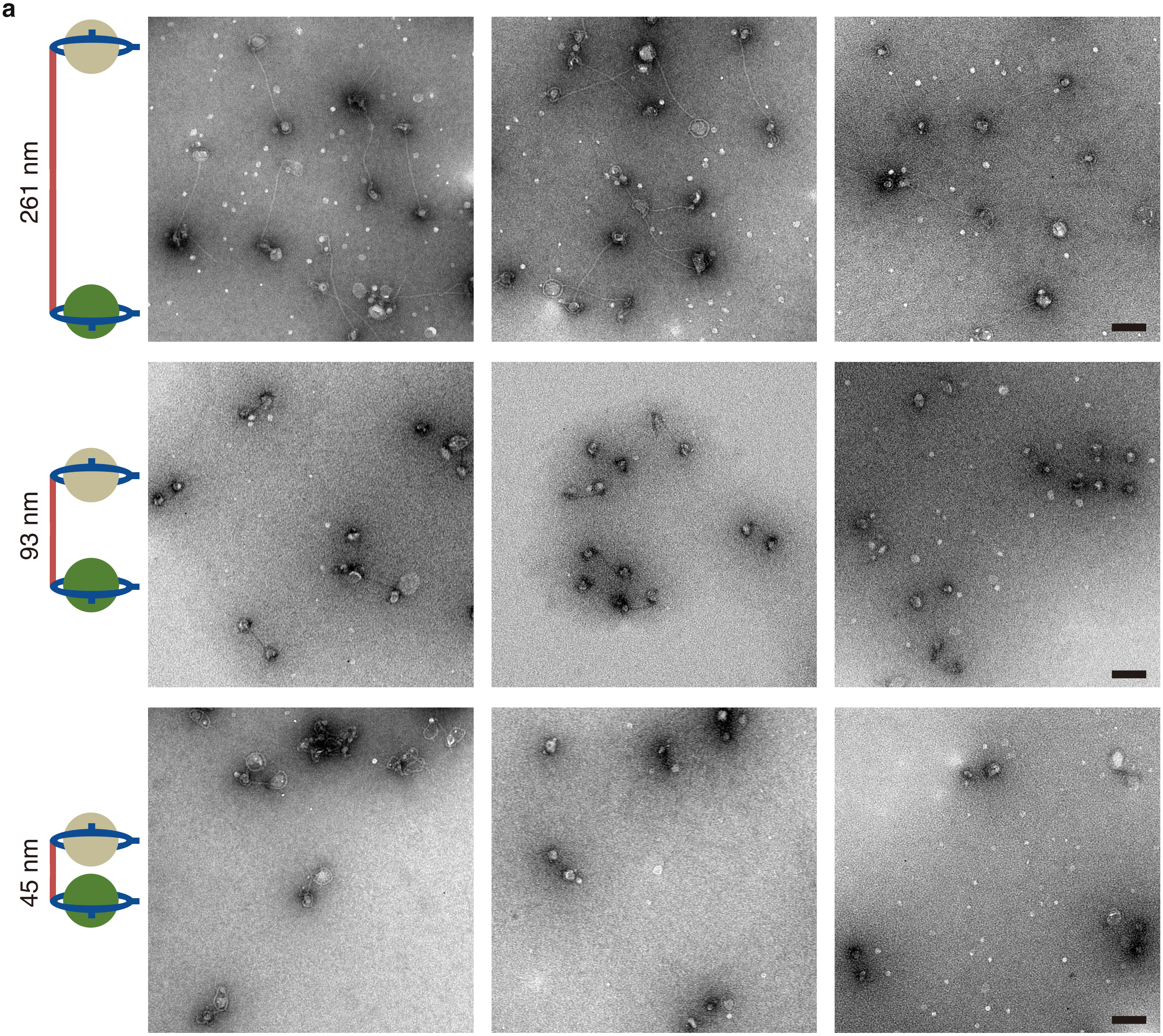


**b**

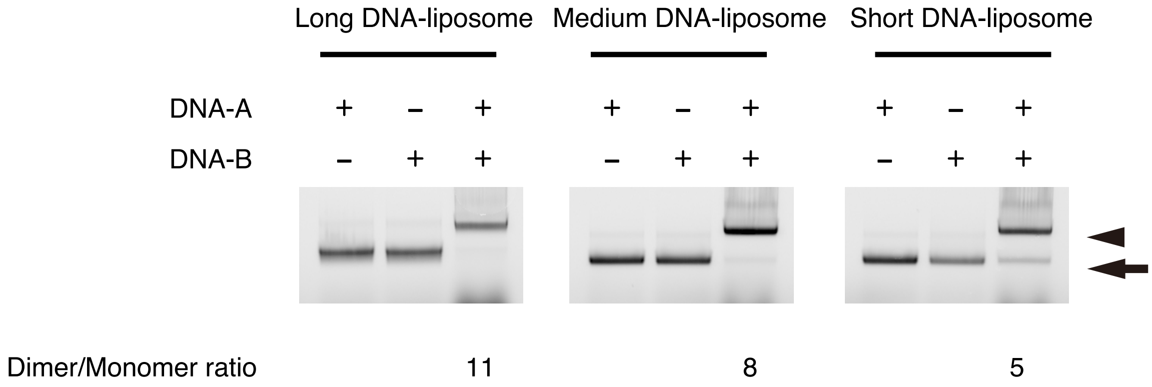


Supplementary Figure 3
